# Evolutionary divergence of LRRK2 interaction domains contributes to human- and mouse-specific protein interaction networks

**DOI:** 10.64898/2026.07.17.738869

**Authors:** Luca Ballotto, Pasquale Miglionico, Yibo Zhao, Luigi Bubacco, Francesco Raimondi, Elisa Greggio, Claudia Manzoni

**Author notes:** These authors contributed equally (Francesco Raimondi, Elisa Greggio, Claudia Manzoni).

## Abstract

Leucine-rich repeat kinase 2 (LRRK2) is a complex multidomain protein whose catalytic and protein–protein interaction domains regulate a wide range of cellular processes. To investigate whether evolutionary divergence of these domains contributes to species-specific differences in LRRK2 biology, we combined phylogenetic, sequence, interactome and structural analyses of human and mouse LRRK2. Phylogenetic analysis revealed that the catalytic core predates the acquisition of the N-terminal and C-terminal protein-protein interaction domains during LRRK2 evolution. Accordingly, despite the high overall sequence similarity between human and mouse LRRK2, sequence divergence was not uniformly distributed across the protein but was concentrated within protein–protein interaction domains, whereas the catalytic ROC–COR– kinase core displayed markedly higher conservation. Consistent with this pattern, comparison of curated human and mouse interactomes revealed substantial differences in protein interaction networks and associated biological pathways. Structural modelling of a subset of interactors further showed that predicted interaction interfaces are enriched for residues that differ between the two species, providing a structural rationale for altered interaction specificity. Together, these findings support the view that evolutionary divergence of LRRK2 protein–protein interaction domains contributes to species-specific interactome organization. These results provide an evolutionary framework for interpreting differences between human and mouse LRRK2 and highlight the importance of considering species-specific interaction networks when translating findings from experimental models.

## Introduction

Mutations in the Leucine-rich repeat kinase 2 (LRRK2) coding gene are the leading cause of monogenic Parkinson’s disease (PD) (Kmiecik et al., 2024; Paisán-Ruíz et al., 2004; Zimprich et al., 2004). LRRK2 is a large, multidomain protein featuring both a GTPase domain (ROC) and a serine/threonine kinase domain (KIN) linked together by the C-terminal of ROC (COR), and 4 protein-protein interaction domains: the N-terminal Armadillo domain (ARM), Ankyrin repeats (ANK) and leucine-rich repeats (LRR), and the C-terminal WD40 domain (Alessi and Pfeffer, 2024; Zhang and Kortholt, 2023; C. Zhu et al., 2023). From a functional point of view, LRRK2 is involved in mediating a remarkable variety of biological processes, including vesicular trafficking through the phosphorylation of Rab GTPases (Steger et al., 2016), cytoskeletal dynamics at the synapse (Tombesi et al., 2025), and at centrosomes (Eckert et al., 2026), mitochondrial dynamics (Saha et al., 2009; Singh et al., 2019), ciliogenesis (Iannotta et al., 2024; Steger et al., 2017; Yeshaw et al., 2023), and inflammation (Ballotto et al., 2026; Wallings and Tansey, 2019). This remarkable functional diversity of LRRK2 is thought to arise from its ability to engage with multiple protein partners within functional networks. Recent network-based analyses of the striatal LRRK2 interactome have identified highly connected protein clusters specifically associated with the development of the midbrain and the organization of the lamellipodium (Zhao et al., 2023). Characterising LRRK2 interactions within protein networks not only provides important insights into its physiological functions, it also helps reveal the molecular perturbations associated with disease states.

Pathogenic LRRK2 mutations, particularly the most common G2019S variant, have been reported to consistently increase the kinase activity of LRRK2 with consequent phosphorylation of a subset of Rab GTPases. Moreover, Rab phosphorylation and LRRK2 autophosphorylation at serine 1292 are increased even in idiopathic PD patients, suggesting a pivotal role of the kinase beyond the limited subset of genetic PD cases (Di Maio et al., 2018). Several mice models have been generated to elucidate the pathogenic mechanisms underlying LRRK2-PD including knock-out, knock-in, and overexpression models as well as bacterial artificial chromosome (BAC) transgenic mice (Li et al., 2009; Melrose et al., 2010; Tong et al., 2010). Strikingly, most of these murine models fail to display dopaminergic neurodegeneration, the fundamental hallmark of PD neuropathology (Domenicale et al., 2023). The only exception to this is for a handful of transgenic lines harboring the human LRRK2 protein with the G2019S mutation (Arbez et al., 2020; Chen et al., 2012; Ramonet et al., 2011; Xiong et al., 2018). However, not all the hG2019S mice display neurodegeneration, which may be due to heterogeneity in the transgene expression level, choice of promoter, or genetic background (Domenicale et al., 2023). Nonetheless, dopaminergic degeneration has never been reported in knock-in mice harboring the G2019S variant in the endogenous murine gene.

Another interesting difference between the human and the murine LRRK2 genes comes from analyses of neuroinflammation. Alongside with neurodegeneration, neuroinflammation is another hallmark of PD which is thought to be amplified by pathogenic LRRK2 mutations (McGeer et al., 1988; Russo et al., 2014). This is also corroborated by the fact that pathogenic LRRK2 mutations, outside of PD, have been linked to chronic inflammatory diseases such as Crohn’s disease (Franke et al., 2010). Coherently, LRRK2 is highly expressed in multiple peripheral immune cells, where it is upregulated by interferon-γ signaling and enhances NF-κB-mediated immune responses (Gardet et al., 2010). Interestingly, humanized mice models harboring the human LRRK2 mutant gene and its promoter, display enhanced LRRK2 upregulation downstream interferon-γ signaling (Beilina et al., 2026). This effect has been shown to be mediated by the presence of STAT1 binding sites specifically within the human LRRK2 promoter (Beilina et al., 2026).

These data point to an important but still unresolved question: are there intrinsic structural and network-level differences between human and murine LRRK2 that impair the ability of mouse models to recapitulate PD pathology? Addressing this gap is particularly important because most knock-in models, despite carrying pathogenic mutations in the endogenous gene, do not develop robust dopaminergic neuronal loss, suggesting that mutation-driven kinase hyperactivation may not be sufficient in the murine context.

To date, few comparative studies have directly assessed functional differences between human and mouse LRRK2. One such study investigated the stability and activity of the two LRRK2 orthologs in HEK293FT cells (Langston et al., 2019). Interestingly, mouse LRRK2 displayed an increased affinity to chaperone proteins Hsp90, Hsc70, and CHIP, which possibly make mouse LRRK2 more stable than its human counterpart (Langston et al., 2019). Moreover, the kinase activity of the human LRRK2 was reported to be higher than mouse LRRK2, demonstrated by higher substrate phosphorylation and autophosphorylation (Langston et al., 2019).

Collectively, these findings point to the existence of potential functional divergence between human and mouse LRRK2, highlighting a complex and multifaceted biological landscape. A systematic comparative analysis of the two proteins could provide valuable insights, both into the limitations of existing mouse models in reproducing key aspects of PD and into the mechanistic links between LRRK2 and disease pathogenesis. Such studies may be especially informative given the apparent lack of a direct association between mouse LRRK2 and disease, in contrast to the well-established role of human LRRK2.

Here, we performed an in-silico comparison of human and mouse LRRK2, focusing on differences in protein sequence conservation, 3D structure, and interaction behaviour. We highlight key differences between the two orthologues which may have important implications for disease modelling.

## Materials and Methods

### Sequence retrieval and domain definition

Human and mouse LRRK2 protein sequences were retrieved from UniProt (human: Q5S007, mouse: Q5S006) and aligned to quantify sequence divergence between orthologues in R using the pwalign and Biostrings packages. LRRK2 domain boundaries were defined based on annotations from InterPro (UniProt entries Q5S006 and Q5S007). Where the domain intervals didn’t match for the two orthologous, the lowest starting position and highest end position were selected to define that domain. The final domain intervals were defined as follows: (17–649), ANK (679–808), LRR (957–1314), ROC (1328–1511), COR (1524–1858), KIN (1879–2146), and WD40 (2165–2509).

For the analysis correlating sequence divergence with structural deviation (RMSD), the LRR domain interval was modified to exclude the N-terminal hinge region (residues 957–984). Preliminary structural alignments indicated that this flexible segment introduced a disproportionate contribution to the overall RMSD despite limited sequence divergence, thereby obscuring the relationship between sequence variation and structural conservation within the core LRR fold. Therefore, RMSD calculations for the LRR domain were performed using residues 985–1305 only. All other analyses were performed using the original InterPro-derived domain boundaries.

Mismatch rates were calculated as the proportion of non-identical amino acids between human and mouse LRRK2 within each domain, normalized by domain length. Results were visualized as bar plots and summarized in a table reporting domain length, number of mismatches, and mismatch percentage.

### MIST LRRK2 interactome dataset

LRRK2 protein–protein interaction (PPI) datasets were downloaded from the Molecular Interaction Search Tool (MIST v5.0; November 2025, https://fgrtools.hms.harvard.edu/MIST/) (Hu et al., 2018). Interactions categorized as “high-confidence” and “medium-confidence” were retained, while “low-confidence” interactions were excluded. This resulted in a shorter list of 394 interactors for human LRRK2 and 84 interactors for mouse LRRK2. Interactor lists were processed in R to remove missing values and duplicated entries.

To enable comparison between species, mouse interactors were converted to their corresponding human orthologues using the g:Profiler tool **g:**Orth (https://biit.cs.ut.ee/gprofiler/orth) (Kolberg et al., 2023). When orthologues were available, mouse gene names were replaced with the corresponding human ortholog symbols. Shared and species-specific interactors were identified, and the overlap between datasets was visualized using a Venn diagram.

### Functional enrichment analysis

Functional enrichment analysis of the human and mouse LRRK2 interactomes was performed using g:Profiler g:GOSt, restricting the analysis to GO Biological Process (GO:BP) terms. The statistical correction method was set to g:SCS, with a significance threshold of adjusted p-value < 0.05. To reduce redundancy and overly broad terms, enrichment results were filtered by term size where applicable, with a cutoff of 50 terms. GO:BP terms were grouped by semantic similarity using keywords into semantic classes. The most significant term in each semantic class was selected to represent the p-value of the entire class (g:SCS adjusted p-value). Semantic classes were further organized into broader functional groups linking multiple classes, which were color-coded for visualization.

Additional pathway enrichment analysis and term clustering were performed using Metascape (Zhou et al., 2019). to validate major functional themes and support interpretability across interactome lists.

### Structural modeling and contact-site analysis

Structural comparison between human and mouse LRRK2 was performed using available predicted structures (AlphaFold) and structural visualization tools. To explore potential binding interfaces between mouse LRRK2 and mouse-specific interactors, protein–protein complex predictions were generated using AlphaFold 3 (Abramson et al., 2024) for selected interactors (Ctnnb1, Axin1, Cdk5r2, Stxbp1 and Vamp2). Predicted contact residues at the LRRK2 interface were extracted and visualized with UCSF ChimeraX (Meng et al., 2023).

To assess whether predicted mouse-specific contact sites corresponded to species-divergent regions, contact residues were compared to amino acid mismatch positions identified in the human–mouse sequence alignment. The overlap between interface residues and mismatches was quantified as the proportion of contact residues located in non-conserved positions.

### Phylogenetic analysis

LRRK2 orthologs were selected from MetaPhOrs (release-201601) (Pryszcz et al., 2011) considering the set of proteins with an orthology confidence score greater than 0.5. We then used the InterPro database (release 82.0) (Blum et al., 2021) to identify the domains of each ortholog and we excluded only proteins with less than two domains in common with the human LRRK2. This allowed us to reduce the size of the tree, improving its interpretability. The multiple sequence alignment (MSA) was built with ClustalOmega (Sievers et al., 2011) using the options --iter=5 --full. The maximum likelihood phylogenetic tree was inferred from the MSA with IQ-TREE (version 1.6.12) (Nguyen et al., 2015) using the built-in ModelFinder function to choose the model of rate heterogeneity across sites. Tree images were drawn using the ete3 package (Huerta-Cepas et al., 2016).

## Results

### Phylogenetic analysis of LRRK2 reveals an ancient origin of its multi-domain architecture

LRRK2 is a complex multidomain protein containing several catalytic and protein-interaction domains. To gain insight into how these domains contribute to the functional specialization of LRRK2, and whether they may account for the species-specific differences observed between the human and mouse proteins, we investigated the evolutionary history of LRRK2’s domain architecture.

We built a phylogenetic tree of LRRK2 orthologs retrieved from MetaPhOrs (Fig. 1). As expected, most vertebrate LRRK2 orthologs displayed the same domain architecture as human LRRK2 (ARM–ANK–LRR–ROC–COR–KIN–WD40). Notably, this same architecture was also present in other deuterostomes (the lancelet *Branchiostoma floridae* and the sea urchin *Strongylocentrotus purpuratus*), in a lophotrochozoan (the oyster *Crassostrea gigas*), and in a cnidarian (the sea anemone *Nematostella vectensis*), indicating that the complete LRRK2 architecture may be considerably more ancient than the vertebrate lineage, consistent with previous evolutionary analyses identifying a bona fide LRRK2 ortholog in cnidarians (Marín, 2008).

**Figure 1:**
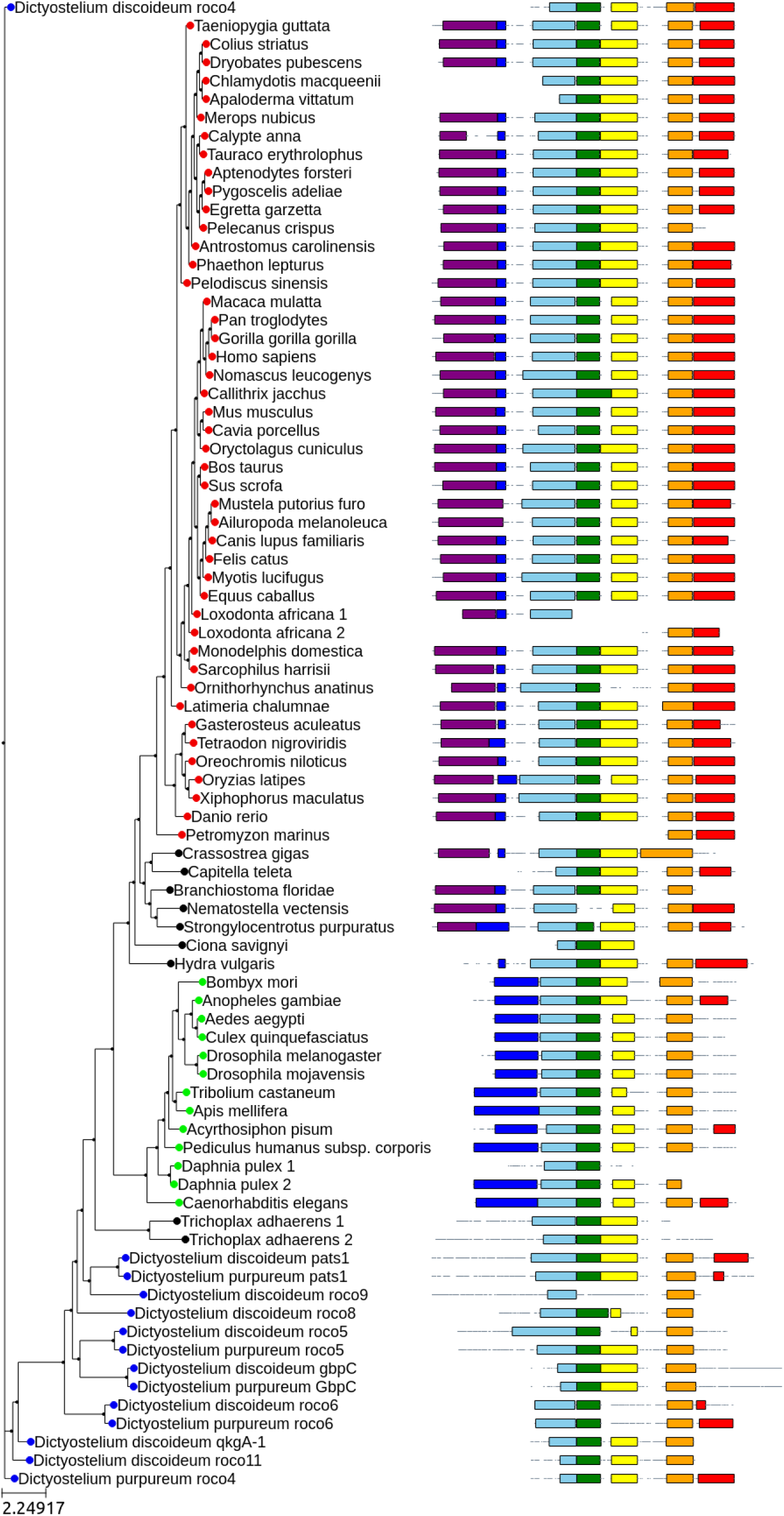
Phylogenetic tree showing the evolution of LRRK2 architecture. Vertebrates are represented with a red dot, Ecdysozoa are represented by a green dot and amoebas of the *Dictyostelium* genus are represented by a blue dot, black dots indicate species which are not included in the previous groups. On the right part of the tree we show the architecture of LRRK2 orthologs. Each domain is shown in a different color: ARM in purple, ANK in blue, LRR in skyblue, ROC in green, COR in yellow, KIN in orange and WD40 in red.

In contrast, LRRK2 orthologs in Ecdysozoa lacked the N-terminal ARM domain and possessed a longer ANK region, an architecture more reminiscent of human LRRK1 (the closest LRRK2 paralog in vertebrates). The best-characterized protein of this group is LRK-1 in *Caenorhabditis elegans*. Because the ARM-containing architecture is found in both cnidarians and lophotrochozoans, the most parsimonious interpretation is that the ARM domain was secondarily lost along the ecdysozoan stem.

The tree also includes two Roco proteins lacking a kinase domain from the placozoan *Trichoplax adhaerens* and several Roco proteins with a kinase domain from *Dictyostelium discoideum* and *Dictyostelium purpureum*. Their placement indicates that the catalytic core (ROC and KIN) together with the COR and LRR domains emerged earliest in Roco evolution, whereas the accessory ARM, ANK, and WD40 domains became fixed within the LRRK2 architecture later, but before the cnidarian–bilaterian last common ancestor. The long-term conservation of these accessory domains across animals, together with their selective loss in specific lineages, is consistent with a role in shaping the functional specialization of LRRK2, and, more broadly, the diversification of ROCO protein functions despite conservation of the catalytic ROC–COR core (Tomkins et al., 2018).

The progressive acquisition of protein–protein interaction domains during LRRK2 evolution led us to hypothesize that these regions may have remained more evolutionarily plastic than the conserved catalytic core, thereby contributing to species-specific specialization of the LRRK2 interactome. To test this hypothesis in a translationally relevant context, we next focused our analysis on the human and mouse orthologues.

### The variability between human and mouse LRRK2 is concentrated in protein-protein interaction domains

Human and mouse LRRK2 share a high degree of sequence homology (Fig. 2A), which is estimated to be approximately 88% (Langston et al., 2016). However, whether this homology is equally distributed across the various protein domains warrants further investigation. We calculated the % of mismatches for each protein domain and found significant variability, which is not homogeneously distributed across the 7 main domains of the protein (Fig. 2B). In particular, the enzymatic core is highly conserved between the orthologues, with a mismatch rate of 4.85-5.98%, while most of the variability is concentrated in the protein-protein interaction (PPI) domains, ranging from 14.6 to 17.5% (Fig. 2C).

**Figure 2:**
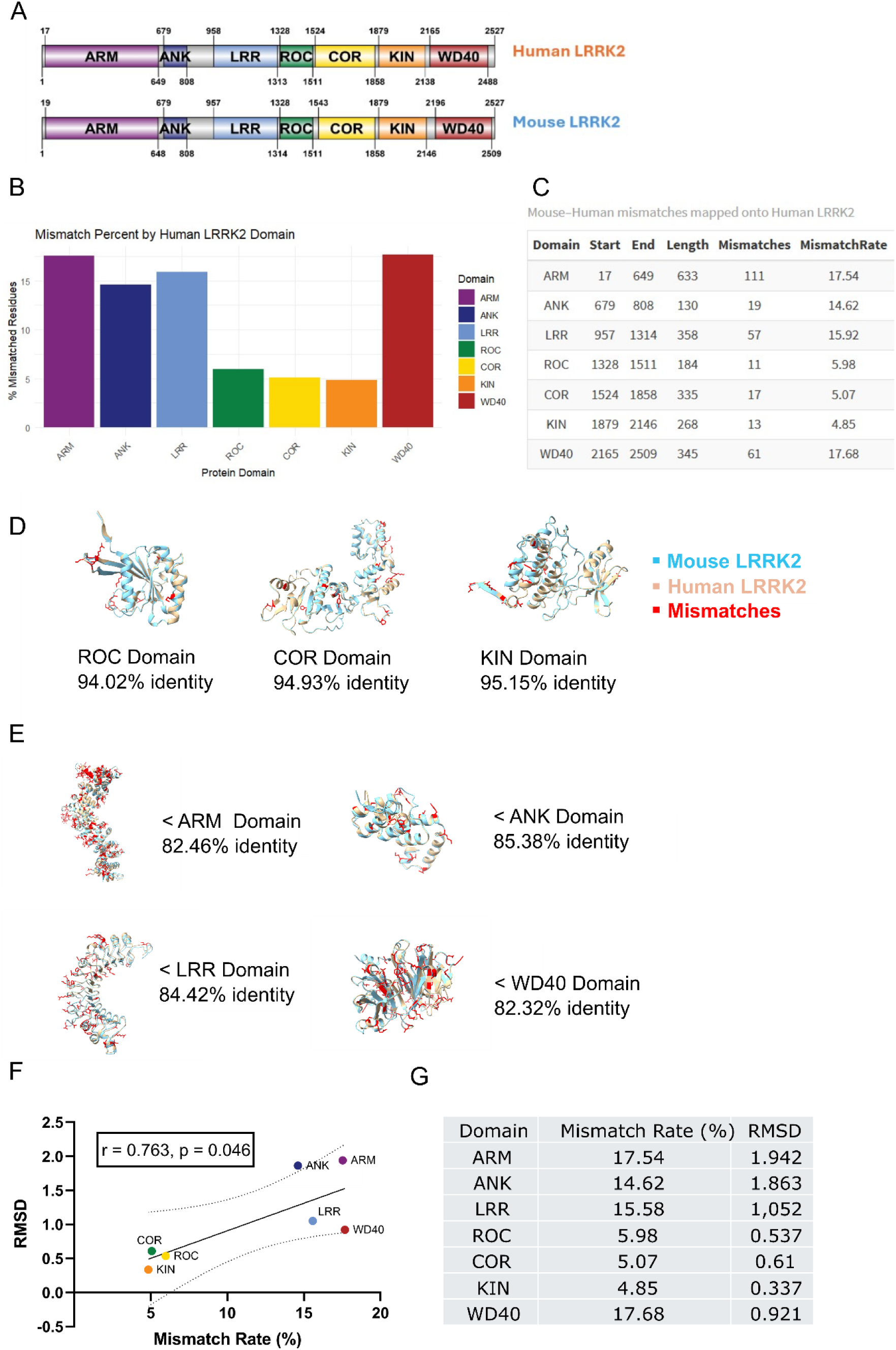
The variability between human and mouse LRRK2 is concentrated in protein-protein interaction domains. (A) Schematic representation of human and mouse LRRK2 domain intervals. (B) Bar plot of the % of mismatches (coding variation) between human and mouse LRRK2, plotted domain by domain. (C) Summary table of coding variation between each protein domain. Mouse–human mismatches were mapped onto human LRRK2 residue coordinates. Due to an insertion/deletion in the ARM domain, residue positions differ between species in this region, whereas downstream domains are aligned without positional offset. (D) 3D reconstruction of murine (light blue) and human (light brown) enzymatic core domains. Mismatched residues are highlighted in red. (E) 3D reconstruction of murine (light blue) and human (light brown) protein-protein interaction domains. Mismatched residues are highlighted in red. (F) Scatter plot showing the correlation between sequence divergence and 3D structural deviation across individual LRRK2 domains. Sequence divergence between human and mouse LRRK2 is quantified as mismatch rate (%), while structural deviation is represented by RMSD (Å) calculated from AlphaFold-predicted structures after structural alignment using ChimeraX MatchMaker. Each point corresponds to a distinct domain. R² = 0.582, r = 0.763, p = 0.046. (G) Summary table of domain-specific sequence mismatch rates (%) and corresponding RMSD (Å) values.

Next, we assessed whether 2D sequence divergence is positively correlated to 3D structural deviation across the individual LRRK2 domains. We overlapped each domain of the two orthologues and quantified RMSD (Å) and % of sequence identity (Fig. 2D,E). Linear regression analysis revealed a significant association between mismatch rate and RMSD (R² = 0.582, P = 0.046), indicating that protein-protein interaction domains, which exhibit greater sequence divergence, tend to also exhibit larger 3D structural differences between human and mouse LRRK2 (Fig. 2F,G).

### Comparative analysis of human and mouse LRRK2 interactomes identifies shared and species-specific binding partners and biological processes

Although the enzymatic core of LRRK2 is highly conserved between human and mouse, the enrichment of mismatches within protein–protein interaction domains suggests that species-specific differences may emerge at the level of binding partners and complex formation. Since LRRK2 function is strongly influenced by its interaction network, we next investigated whether the reported LRRK2 interactome differs between species. To test this hypothesis, we downloaded a dataset of protein-protein interactions from MIST and checked which PPI were reported for either human LRRK2, mouse LRRK2 or both. In total, the dataset included 394 high-confidence interactions for human LRRK2 and 84 for mouse LRRK2 (Supplementary Table 1). Among these, 79 interactors are shared, while 315 are reported only for human LRRK2 and 5 only for mouse LRRK2 (Fig. 3A). Although the human LRRK2 interactome is substantially larger than the mouse interactome—likely owing to the stronger focus of the field on human LRRK2—it was nonetheless striking to identify five proteins that were uniquely associated with the murine protein. To assess whether these interactions were simply excluded by our confidence threshold, we also examined the complete MIST dataset including low-confidence interactions. Of the five mouse-specific interactors (Ctnnb1, Axin1, Cdk5rap2, Stxbp1, Vamp2), only Ctnnb1 was reported as a low-confidence human LRRK2 interactor, whereas the remaining four were not detected in the human dataset (Supplementary Table 2). This finding further supports the existence of species-specific features within the LRRK2 interaction network.

**Figure 3:**
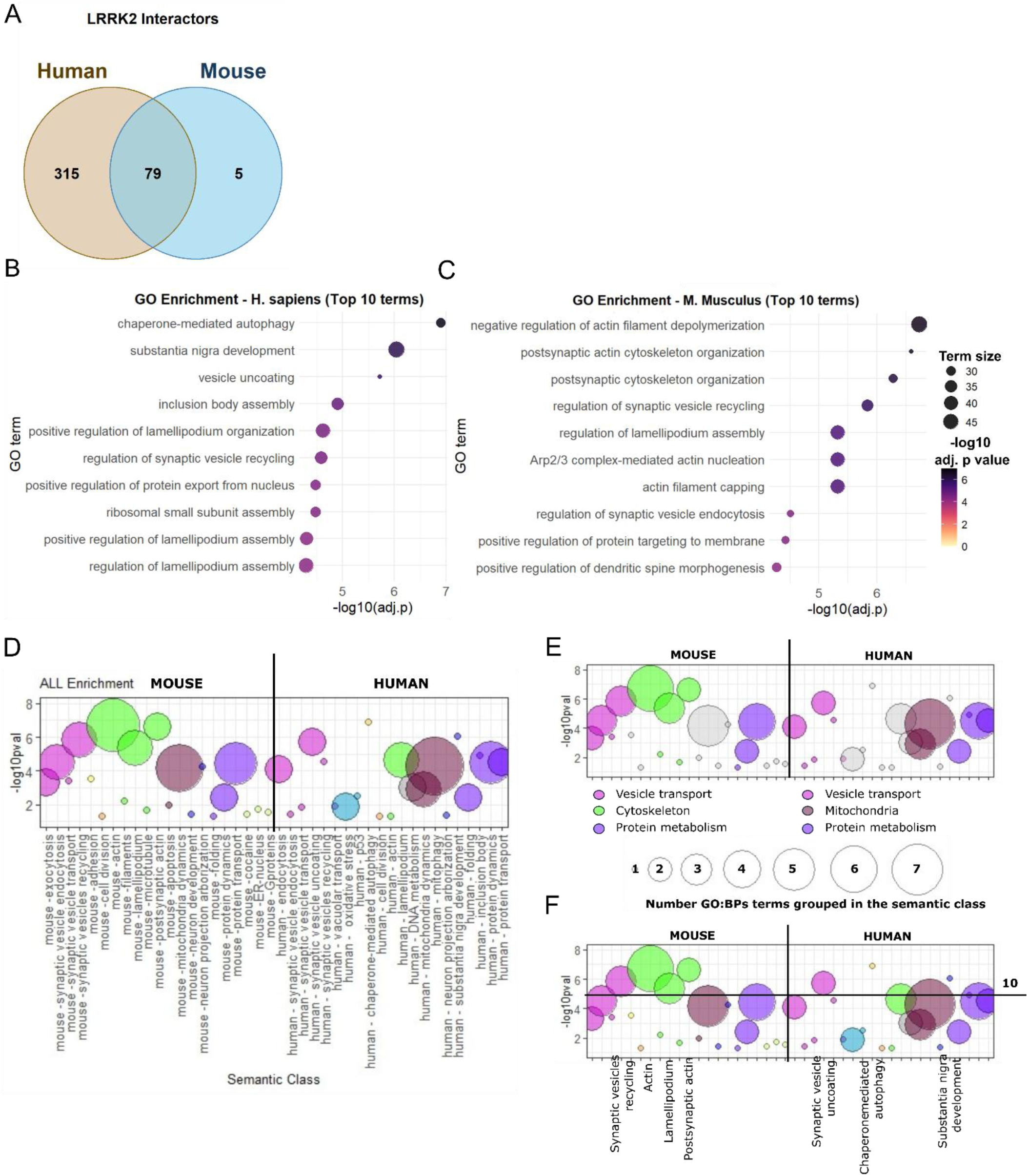
Comparison of Human and Mouse LRRK2 interactome functional enrichment. (A) Venn diagram of shared and unique LRRK2 interactors. (B) Functional enrichment bubble plot of human LRRK2 interactome (C) Functional enrichment bubble plot of mouse LRRK2 interactome. Dot color is proportional to −log10(adjusted p value), and dot size is proportional to term size. (D) GO:BP terms grouped by semantic similarity using keywords into semantic classes. The most significant term in each semantic class was selected to identify the p-value of the entire class (adjusted p-value). The bubble size is directly proportional to the number of GO:BPs grouped in the semantic class. The semantic classes have the same color if they belong to the same functional group. (E) Mouse and human enrichment plotted together for comparison, highlighting the functional groups that collect the majority of the GO:BP terms. (F) Mouse and human enrichment plotted together for comparison highlighted the semantic classes with adjusted p-value < 10^-5^.

We next performed Gene Ontology functional enrichment with gProfiler to assess the overall functional properties of the two interactomes (Fig. 3B,C, Supplementary Table 3). After term size thresholding, the top 3 most significant GO:BPs describing the human LRRK2 interactome (based on enrichment p value) were: “chaperone-mediated autophagy”, “substantia nigra development” and “vesicle uncoating” (Fig. 3B) and were not present in the mouse interactome functional enrichment.

The enrichment of chaperone-mediated autophagy (CMA) was driven by the human-specific LRRK2 interactors HSPA8, LAMP2, BAG3, EEF1A1, and EEF1A2. HSPA8 (Hsc70) and LAMP2A constitute the core machinery of CMA (Orenstein and Cuervo, 2010), whereas BAG3 functions in chaperone-assisted protein quality control and autophagic responses (Minoia et al., 2014). (Albanese et al., 2019; Orenstein et al., 2013) Given the well-established role of LRRK2 in autophagy, the enrichment of CMA-associated interactors was not surprising. Notably, this emerged as a human-specific feature in our analysis, suggesting that human-specific components of the LRRK2 interactome may contribute to CMA-mediated proteostasis mechanisms relevant to PD. These findings raise the possibility that such interactions might be less prominent in the mouse protein, highlighting potential species-specific differences in LRRK2 function.

The top 3 hits for the mouse interactome were “negative regulation of actin filament depolarization”, “postsynaptic actin cytoskeleton organization” and “regulation of synaptic vesicle recycling”. While the first two terms were missing in the human enrichment, the enrichment of synaptic vesicle recycling pathways is shared by both interactomes, suggesting that synaptic vesicle dynamics and, more broadly, endocytic processes are represented in both the human and murine LRRK2 interactomes. Interestingly, cytoskeletal-related terms are considerably more prominent in the mouse enrichment analysis than in the human dataset (13/42 terms in the mouse enrichment, 4/40 terms in human enrichment, Supplementary Table 3), highlighting a potential functional divergence between the two species and suggesting that LRRK2 may engage more extensively with cytoskeletal processes in the murine context.

Similarly, additional analysis with Metascape highlighted cytoskeleton-related biological processes as more related to the murine LRRK2 interactome in comparison with the human LRRK2 interactome (Sup. Fig. 1).

Because the initial analysis focused exclusively on the top enriched terms, it may not have fully captured the overall structure of the enrichment results. We therefore performed a complementary global analysis in which enriched GO terms were semantically grouped and prioritized based on their statistical significance and the proportion of terms within each semantic cluster. Categories with stronger statistical support and larger term representation were highlighted as the dominant functional themes within the dataset.

Interestingly, only the “synaptic vesicle” class was shared between the two interactomes. In contrast, the classes “actin,” “lamellipodium,” and “postsynaptic actin” were selectively enriched in the mouse interactome, whereas “chaperone-mediated autophagy” and “substantia nigra development” were specifically enriched in the human interactome. Collectively, these results confirm the species-specific differences in the functional profiles of LRRK2 interactomes already highlighted by the top enriched terms.

### Structural divergence in LRRK2 supports species-specific protein–protein interactions

The interactome analysis revealed a markedly larger number of human-specific interactors compared to mouse-specific interactors, and functional enrichment suggested distinct biological processes across the two networks. To test whether mouse-specific interactions could be structurally supported by mouse-exclusive sequence features, we next performed structural predictions to identify binding interfaces between mouse LRRK2 and the 5 specific mouse-specific interactors (Ctnnb1, Axin1, Cdk5rap2, Stxbp1, Vamp2) and quantified how these contact sites differ between human and mouse LRRK2 (Fig. 4A). Structural analysis revealed that the ARM domain of mouse LRRK2 is probably mediating the interaction with the mouse-specific interactors, and the WD40 is also involved in the binding with Axin1 (Fig 4B). Interestingly the ARM and WD40 domains were already identified as those with the largest mismatch between human and mouse LRRK2; in this specific case, and focusing on the aa predicted to be involved in the binding, 5 to 17% of each interactor’s contact sites are divergent from human LRRK2, which may contribute to differences in the interaction specificity (Fig. 4C).

**Figure 4:**
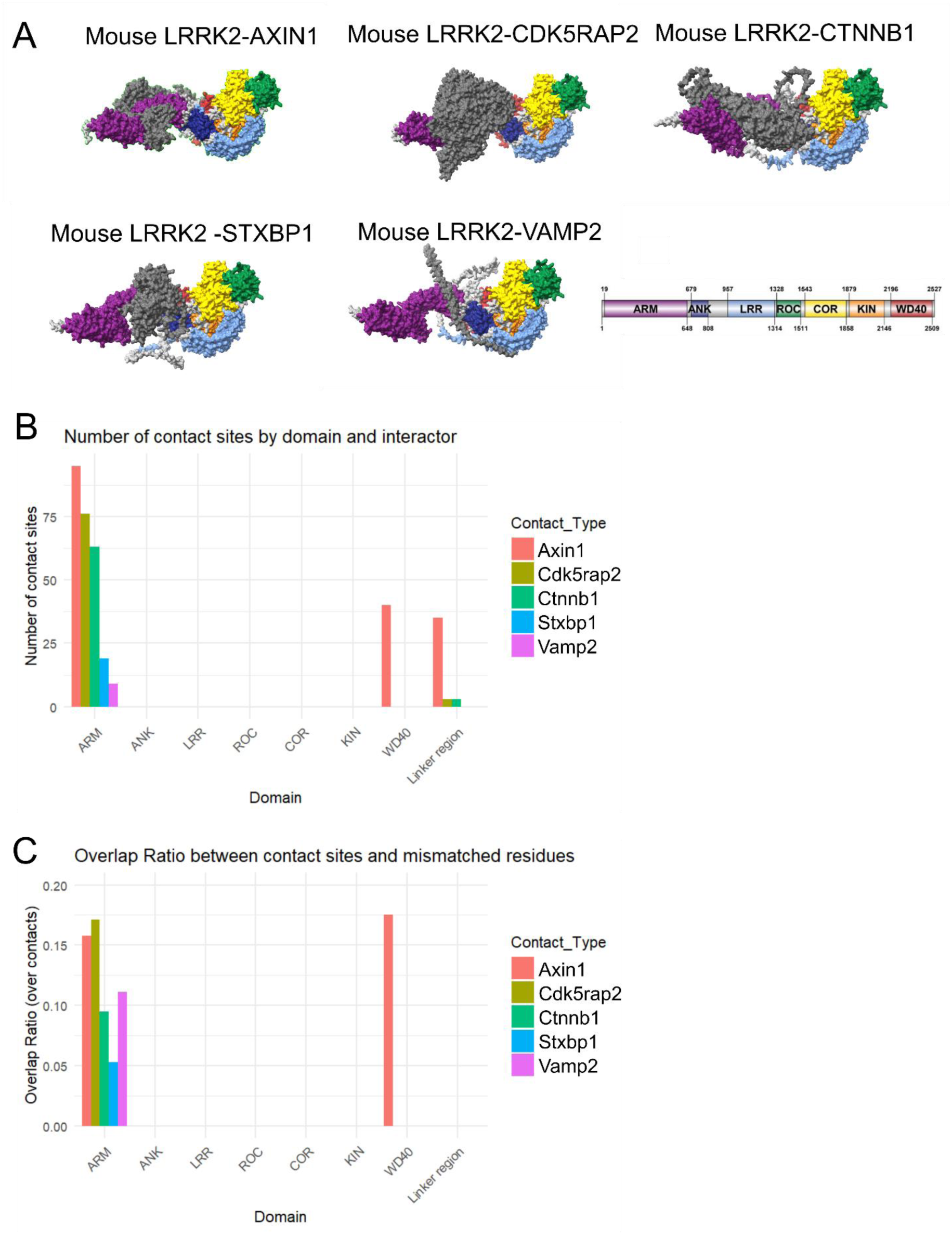
analysis of contact sites between mouse LRRK2 and mouse-specific interactors. (A) AlphaFold 3 predictions of mouse LRRK2 binding to Axin1, Cdk5rap2, Ctnnb1, Stxbp1 and Vamp2. LRRK2 domains color scheme is shown, interactors are colored in dark gray (B) Quantification of number of contact sites between mouse-specific interactors and each LRRK2 domain (C) Fraction of mouse LRRK2 contact-site residues that are mismatched relative to human LRRK2, calculated as mismatched contact-site residues divided by total contact-site residues for each domain.

### Conservation of Rab-binding interfaces across species

Next, we assessed whether the contact sites of shared interactors between human and mouse LRRK2 are conserved across species. To do this, we performed the same structural analysis on a subset of Rab GTPases (Rab10, Rab29, and Rab8A), which are bona-fide LRRK2 interactors shared between mouse and human interactomes. Structural predictions of mouse LRRK2 in complex with Rab proteins revealed that, similarly to mouse-specific interactors, binding is primarily mediated by the ARM domain for Rab10 and Rab29, whereas Rab8A exhibits a broader interaction profile involving additional regions, including COR, KIN, and linker regions (Fig. 5A–B).

**Figure 5:**
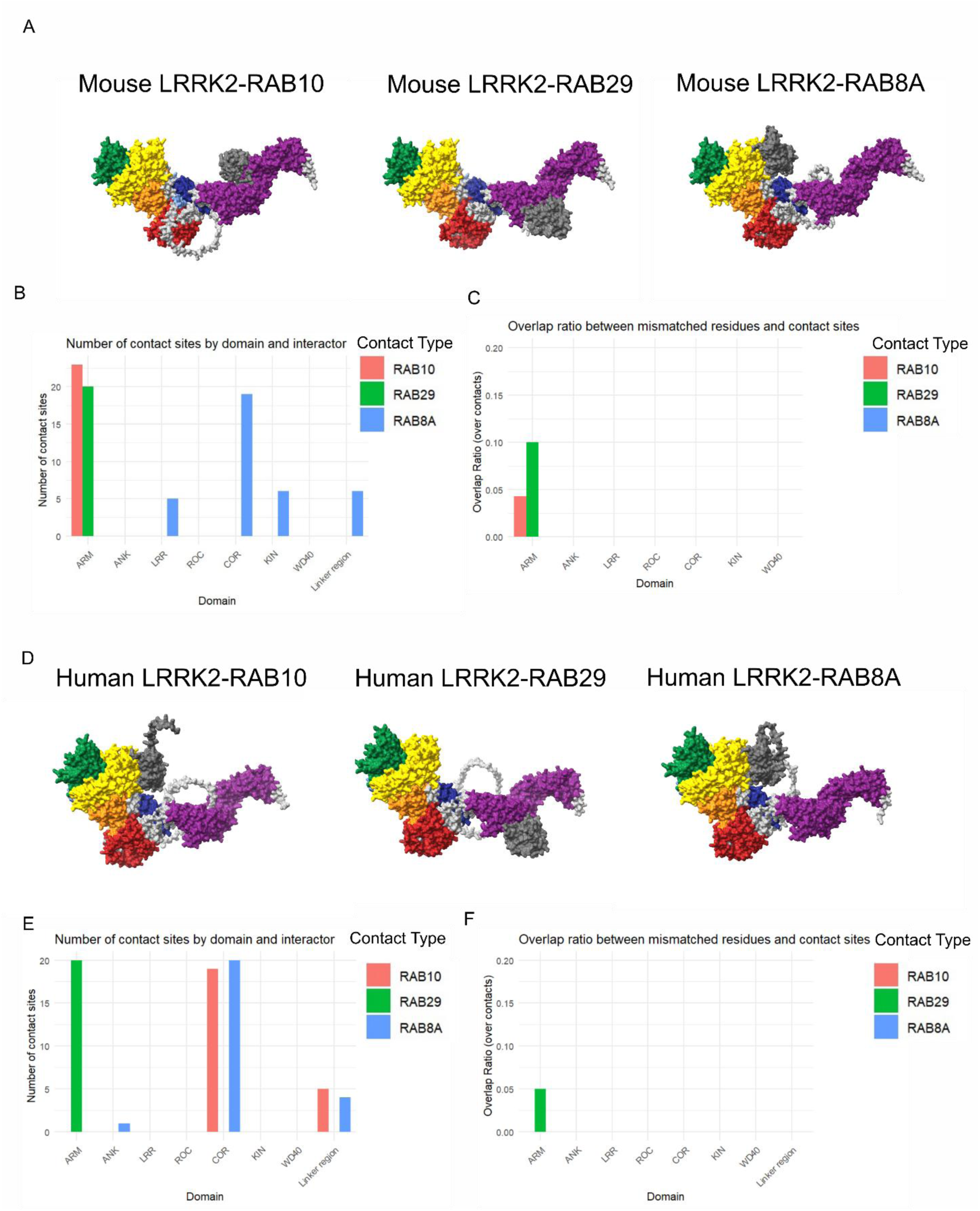
reduced sequence variability at Rab-Binding contact sites between mouse and human LRRK2. (A) Structural models of mouse LRRK2 in complex with different mouse Rab GTPases (Rab10, Rab29, and Rab8A), generated using AlphaFold predictions. LRRK2 domains are color-coded, and interacting Rab proteins are shown in grey. (B) Quantification of contact sites between mouse LRRK2 domains and mouse Rab interactors. The number of contact residues is shown for each domain across Rab10, Rab29, and Rab8A. Interactions are predominantly localized to the ARM domain for Rab10 and Rab29, whereas Rab8A exhibits a broader interaction profile, including contacts within COR, kinase (KIN), and linker regions, indicating differential binding patterns among Rab proteins. (C) Fraction of mouse LRRK2 - mouse Rabs contact-site residues that are mismatched relative to human LRRK2, calculated as mismatched contact-site residues divided by total contact-site residues for each domain. (D) Structural models of human LRRK2 in complex with different human Rab GTPases (RAB10, RAB29, and RAB8A), generated using AlphaFold predictions. LRRK2 domains are color-coded, and interacting Rab proteins are shown in grey. (E) Quantification of contact sites between human LRRK2 domains and human Rab interactors. The number of contact residues is shown for each domain across RAB10, RAB29, and RAB8A. In contrast to mouse LRRK2, human LRRK2 is predicted to interact with RAB10 at the COR domain, similar to RAB8A, while RAB29 interaction residues are predominantly localized to the ARM domain. (F) Fraction of human LRRK2 - human Rabs contact-site residues that are mismatched relative to mouse LRRK2, calculated as mismatched contact-site residues divided by total contact-site residues for each domain.

In contrast to mouse-specific interactors, Rab-binding interfaces showed markedly lower sequence divergence between species. Only a small fraction of contact residues were mismatched between mouse and human LRRK2, with approximately 10% of residues differing in the Rab29 interface and 5% for Rab10. Notably, the interaction interface with Rab8A was fully conserved, with no mismatched residues detected at contact sites (Fig. 5C).

Collectively, these results indicate that conserved interactors such as Rab GTPases engage structurally preserved binding interfaces in LRRK2, whereas mouse-specific interactors preferentially target regions enriched in species-specific sequence variation. This supports a model in which sequence divergence in LRRK2 contributes selectively to species-specific protein–protein interactions while maintaining conserved functional interactions.

We next repeated this analysis by modeling human LRRK2 in complex with human Rab GTPases (RAB10, RAB29, and RAB8A) and assessing the conservation of contact sites relative to mouse LRRK2 (Fig. 5D). Consistent with the mouse models, Rab-binding interfaces were largely localized to the ARM and COR domains, with RAB29 interactions predominantly occurring within the ARM domain, while RAB8A engaged both COR and adjacent regions (Fig. 5E). However, in contrast to mouse LRRK2, the predicted binding interface for RAB10 in human LRRK2 was shifted toward the COR domain, indicating a potential difference in binding mode despite sequence conservation.

Analysis of sequence variability at contact sites revealed minimal divergence between species, with mismatches detected only in the ARM domain. Approximately 5% of contact residues in the human LRRK2–RAB29 interface were mismatched relative to mouse LRRK2 (1/20), while no mismatches were observed for RAB10 or RAB8A contact sites, indicating that these interaction interfaces are fully conserved (Fig. 5F).

Together, these results indicate that Rab-binding interfaces in LRRK2 are highly conserved at the sequence level, but may exhibit species-dependent differences in binding configuration, as suggested by the altered RAB10 interaction site.

## Discussion

Mice are still the most common model system in PD research and have provided invaluable insights on the molecular mechanisms underlying the disease. However, the extent to which murine models can fully recapitulate a human-specific disease is controversial and requires careful investigation (Chang et al., 2022). The aim of this work was to assess whether structural and interactome-level differences between human and mouse LRRK2 could contribute to the limited ability of most murine LRRK2 models to recapitulate key features of PD pathology, particularly nigrostriatal degeneration. Structural and biochemical studies have shown that the enzymatic core of LRRK2, is highly conserved, while less structural information is available for the large N-terminal and C-terminal domains, which primarily mediate protein–protein interactions and regulatory function (Zhang and Kortholt, 2023). Although human and mouse LRRK2 share high overall sequence similarity, our domain-level analysis revealed that sequence divergence is not uniformly distributed across the protein domains. Instead, variability was concentrated within protein–protein interaction domains, while the enzymatic core displayed markedly higher conservation. This pattern is consistent with stronger evolutionary constraints acting on the catalytic machinery, whose functions are likely maintained across species, whereas the surrounding interaction domains appear to be more evolutionarily malleable, allowing lineage-specific tuning of scaffolding, localization and protein–protein interaction networks. The higher frequency of amino acid mismatches within these interaction domains suggest that species differences may primarily influence LRRK2 function through altered interaction networks rather than through major changes in catalytic machinery.

Consistent with this hypothesis, the comparison of human and mouse LRRK2 interactomes revealed substantial asymmetry. The human interactome contained many more reported high-confidence interactors than the mouse interactome, with the majority of interactions appearing human-specific. However, an important limitation to be addressed concerns knowledge bias affecting interactome analysis. Indeed, most interactomics studies involving LRRK2 have been performed in human cell lines, implying that some of the interactions classified as human-specific may simply reflect a lack of systematic testing in murine model systems (Porras et al., 2015). In addition, the smaller size of the mouse interactome dataset may also reflect technical constraints, such as reduced coverage or lower expression in experimental systems.

Therefore, the 315 “human-specific” interactors should not be interpreted as definitive evidence of absence in mouse. Nevertheless, this bias strengthens the potential relevance of the small subset of mouse-specific interactors, as these interactions were detected despite a lower overall density of studies in murine contexts and, with the exception of Ctnnb1, were not replicated in human to the best of our knowledge. Interestingly, a recent BioID screen detected AXIN1 as a proximal interactor of LRRK2 in HEK293 cells (Eckert et al., 2026). This apparent discrepancy likely reflects differences in the underlying experimental approaches. Whereas the affinity purification-based methods that predominate in MIST preferentially detect stable protein complexes, proximity-labelling approaches identify proteins within the local molecular environment, including transient or low-affinity interactors. Accordingly, one possible explanation is that the LRRK2–AXIN1 interaction is sufficiently stable to be detected by affinity purification in mouse, whereas in human it may be weaker or more transient and therefore detectable only by proximity labeling. Taken together, these observations support the hypothesis that human and murine LRRK2 operate within partially distinct protein interaction networks. Such differences in interactome composition may be key determinants of species-specific LRRK2 functions and could help explain why certain aspects of LRRK2 biology and pathology are not fully recapitulated in mouse models.

Functional enrichment suggested that the human LRRK2 interactome is associated with pathways such as chaperone-mediated autophagy, vesicle-related processes, and substantia nigra development, while the mouse interactome showed enrichment for actin cytoskeleton and synaptic-related processes. Together, these results suggest that, even when LRRK2 orthologues share high catalytic conservation, differences in protein interaction networks may lead to divergent biological outputs across species. Interestingly, synaptic defects have been reported in multiple mutant mice models of LRRK2-PD with mutations in the endogenous murine gene (Gupta et al., 2024; Marte et al., 2019; Masotti et al., 2025; Parisiadou et al., 2014; Tombesi et al., 2025), while studies with transgenic mice harboring human LRRK2 did not detect any significant differences in cytoskeletal dynamics (Garcia-Miralles et al., 2015).

To further investigate whether mouse-specific interactions could be supported by structural determinants, we predicted binding interfaces between mouse LRRK2 and selected mouse-specific interactors using AlphaFold 3. These predictions highlighted that contact residues are enriched in regions that overlap with amino acid mismatches relative to human LRRK2. While computational predictions cannot replace experimental validation, these results provide a structural rationale for the possibility that species-divergent residues, particularly within protein– protein interaction domains, may modulate binding affinity or interaction specificity.

It is also important to consider the limitations of machine learning-based structural prediction methods such as AlphaFold. While these approaches have significantly improved the accuracy of single-protein structure prediction, their performance in modelling multi-protein complexes, transient interactions, and dynamic conformational changes remains limited. This is particularly relevant considering that LRRK2 can assemble into homotetramers when binding to certain interactors, such as Rab29 (H. Zhu et al., 2023). Moreover, cryo-EM studies have shown that LRRK2 can adopt either an open, active conformation or a closed conformation, which influences the binding of substrates and modulators (H. Zhu et al., 2023). It therefore remains unclear how faithfully current predictive models such as AlphaFold can capture these dynamic conformational states and the multiple modes of protein–protein interaction associated with LRRK2.

From a translational perspective, these findings contribute to a broader interpretation of why mouse knock-in models carrying pathogenic LRRK2 variants often fail to develop neurodegeneration or other pathological hallmarks. Although mutant LRRK2 can increase kinase activity in both species, the downstream biological consequences may depend on the availability and affinity of protein partners and regulatory networks. Therefore, phenotypic discrepancies between human and murine LRRK2 models may reflect, at least in part, a divergence in interaction landscapes and cellular context between mouse and human.

Another layer of complexity which could explain these discrepancies is the inter-species variability between regulatory regions of LRRK2. In particular, a recent study highlighted an increased induction of human LRRK2 expression compared to mice in response to interferon-γ (IFN-γ), which was explained by the presence of regulatory regions in the human promoter which are absent in the murine counterpart (Beilina et al., 2026). This mechanism is particularly relevant for immune responses in the central nervous system, and it is therefore possible that IFN-γ-mediated responses cannot be properly reproduced in murine models of PD.

In conclusion, this work supports the view that species-specific differences in LRRK2 protein– protein interaction domains may contribute to divergence in interactome composition and functional pathway associations, potentially influencing the translational relevance of murine models. Together, these findings highlight the importance of complementing conventional mouse models with humanized models and human induced pluripotent stem cell (iPSC)-derived systems, which may better recapitulate the molecular context in which human LRRK2 functions.

## Acknowledgements

This work was supported by the European Union – NextGenerationEU (NRRP, Mission 4, Component 2, Investment 1.1 – PRIN, Project 20222LRHCW, CUP C53D23005560006) to EG, and by funding from the associations Associazione Italiana Giovani Parkinsoniani, Prendiamoci Per Mano Aps, and Associazione Turistica Pro Loco Costigliole Saluzzo, as well as by generous individual donations for which we sincerely thank all donors.

