## Supplementary Figure 1 for "Evolutionary divergence of LRRK2 interaction domains contributes to human- and mouse-specific protein interaction networks"

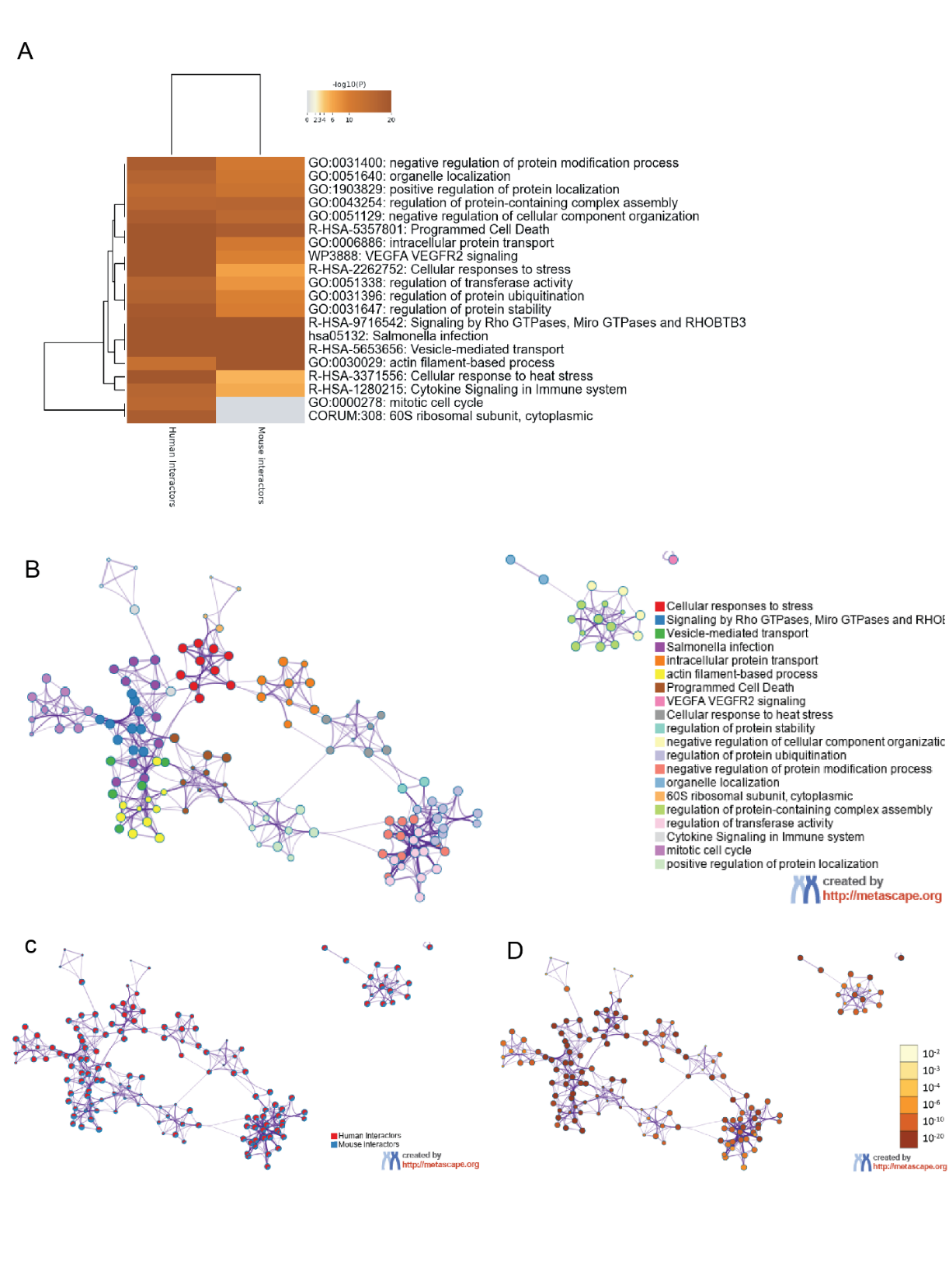

**Supplementary figure 1: Metascape functional enrichment of murine and human LRRK2 interactome.** (A) Top 20 Enriched terms from the functional enrichment analysis, color coded p-value (B) Network layout of functionally enriched terms. Each term is represented by a circle node, where its size is proportional to the number of input genes that fall under that term, and its color represents its cluster identity. (C) The same enrichment network has its nodes displayed as pies. Each pie sector is proportional to the number of hits originated from a gene list. (D) The same enrichment network has its nodes colored by p-value, as shown in the legend.
